# JUMP: replicability analysis of high-throughput experiments with applications to spatial transcriptomic studies

**DOI:** 10.1101/2023.02.13.528417

**Authors:** Pengfei Lyu, Yan Li, Xiaoquan Wen, Hongyuan Cao

**Author notes:** The first two authors should be regarded as Joint First Authors.

## Abstract

**Motivation:** Replicability is the cornerstone of scientific research. The current statistical method for high-dimensional replicability analysis either cannot control the false discovery rate (FDR) or is too conservative.

**Results:** We propose a statistical method, JUMP, for the high-dimensional replicability analysis of two studies. The input is a high dimensional paired sequence of *p*-values from two studies and the test statistic is the maximum of *p*-values of the pair. JUMP uses four states of the *p*-value pairs to indicate whether they are null or non-null. Conditional on the hidden states, JUMP computes the cumulative distribution function of the maximum of *p*-values for each state to conservatively approximate the probability of rejection under the composite null of replicability. JUMP estimates unknown parameters and uses a step-up procedure to control FDR. By incorporating different states of composite null, JUMP achieves a substantial power gain over existing methods while controlling the FDR. Analyzing two pairs of spatially resolved transcriptomic datasets, JUMP makes biological discoveries that otherwise cannot be obtained by using existing methods.

**Availability:** An R package JUMP implementing the JUMP method is available on CRAN (https://CRAN.R-project.org/package=JUMP).

## 1 Introduction

Replicability is the cornerstone of modern scientific research. We study conceptual replicability, where consistent results are obtained from studies using different procedures and populations that target the same scientific questions. Replicability is related to but different from meta-analysis. Both approaches look at cross-experiment summaries. In a meta-analysis, the null hypothesis is that there is no effect in all studies. On the other hand, in replicability analysis, the alternative hypothesis is that the effects exist in all studies and the null hypothesis is a composite null, i.e., at least one study does not have an effect. Methods designed for meta-analysis, such as Fisher’s method (Fisher, 1925), the Šidák’s method (Šidák, 1967), the Lancaster’s method (Lancaster, 1961) and the minimum of *p*-values are not applicable for replicability analysis.

We focus on the replicability analysis of genomic data produced by high-throughput experiments (Li et al., 2011; Philtron et al., 2018; Hung and Fithian, 2020; Bogomolov and Heller, 2022). In a high-throughput experiment, many candidates are evaluated for their association with a biological feature of interest, and those with significant associations are identified for further analysis. We aim to simultaneously identify replicable features from high-throughput experiments in multiple studies. To analyze a single high-throughput experiment, an acute problem is the multiple comparison. A classic method for multiple comparisons is the false discovery rate (FDR) control approach proposed in Benjamini and Hochberg (1995) (BH).

The false discovery rate is defined as the expected value of the ratio of false rejections over total rejections. Suppose we have *m* hypotheses. The BH procedure works as follows. First, order *p*-values *p*_(1)_ *≤* … *≤ p*_(*m*)_. Second, find the largest *i*_0_ such that 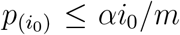. Third, reject hypotheses corresponding to *p*_(1)_, …, *p*_(*i*_0_)_. Under the assumption that *p*-values from the null are independent and follow standard uniform distribution, FDR is controlled at level *απ*_0_, where *π*_0_ is the proportion of null hypotheses. The BH procedure is robust under positive dependence among *p*-values under the null (Benjamini and Yekutieli, 2001).

When we have two studies, an *ad hoc* approach for replicability analysis is to first compute *p*-values for each study. Multiple comparison methods such as BH can be used to claim significance in each study. Replicable findings are those that are significant in both studies. This approach cannot control FDR (Bogomolov and Heller, 2013). Intuitively speaking, if there is no danger that a multiple testing procedure produces false positives, then this *ad hoc* approach would work. However, multiple testing procedures have a non-zero probability of making false positives unless the procedure does not reject. Thus, an approach that provides control over false positives in each study separately does not guarantee control of overall false positives to test replicability (Bogomolov and Heller, 2013). The partial conjunction approach in Benjamini et al. (2009) suggests applying a multiple testing procedure, such as BH, on the maximum of *p*-values across two studies. This provides effective FDR control yet the power is low.

In this paper, we propose a joint super-<u>u</u>niform <u>m</u>aximum *<u>p</u>*-value (JUMP) method for high-dimensional replicability analysis. The null hypothesis consists of three states. The original maximum of *p*-values does not incorporate different states of the null and has a cumulative distribution function far smaller than that of a standard uniform distribution, causing power loss. We use four states for the *p*-value pairs indicating whether they are null or non-null (Chung et al., 2014). Conditional on the hidden states, we compute the cumulative distribution function of the maximum of *p*-values for each state separately. Combining with the proportion estimation for each state, we get a conservative approximation of the cumulative distribution function of the maximum of *p*-values under replicability null. To estimate the proportion of each state, we extend the proportion of null estimation in Storey et al. (2004) from the two-group model to a four-group model. We require *p*-values in each sequence to follow a standard uniform distribution under the null. This assumption is needed for the BH procedure and Storey et al. (2004)’s method of the proportion of null estimation. A step-up procedure is developed to control FDR. By incorporating the composite null feature of replicability analysis, we achieve a substantial power gain. The computation is scalable with a computational cost similar to that of BH.

As proof of concept, we apply JUMP to the replicability analysis of spatial transcriptomic studies. Spatially resolved transcriptomics (SRT) links the transcriptomes to their cellular locations, providing a comprehensive understanding of gene expression profiles in the spatial context of biological systems (Kleino et al., 2022). An important first step toward characterizing the spatial transcriptomic landscape of complex tissues is identifying replicable spatially variable genes (SVGs), genes that have clustered signals in the 2-dimensional space for spatial transcriptomic data. For each study, we can apply existing SVG detection methods to get *p*-values (Edsgärd et al., 2018; Svensson et al., 2018; Sun et al., 2020; Zhu et al., 2021; Hu et al., 2021). The input to our method is a paired *p*-value sequence collected from different tissue sections: mouse olfactory bulb data measured with ST technology (Ståhl et al., 2016) and 10X Visium technology; mouse cerebellum data measured with Slide-seq technology (Rodriques et al., 2019) and Slide-seqV2 technology (Stickels et al., 2021). We show that at the same FDR level, JUMP identifies important replicable SVGs that would otherwise be missed by using existing methods.

## 2 Methods

### 2.1 Background and notations

Suppose we have *m* genes common to two studies. We are interested in identifying genes that display replicable expression patterns. The input of the replicability analysis is a pair of *p*-values from two studies (*p*_1*i*_, *p*_2*i*_), *i* = 1, …, *m*. The null hypothesis for *i*th gene states that it does not show any replicable expression pattern. Let *θ*_*ji*_ denote underlying state of *i*th gene in study *j* (*j* = 1, 2), where *θ*_*ji*_ = 1 indicates *i*th gene is significant in study *j* and *θ*_*ji*_ = 0 otherwise. A four-group model is assumed for the paired *p*-value sequence, i.e.,

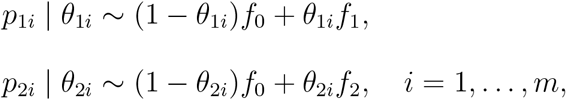

where *f*_0_ is the density function of *p*-values under the null, and *f*_1_ and *f*_2_ denote the nonnull density functions of *p*-values from study 1 and study 2, respectively. Let *τ*_*i*_ = (*θ*_1*i*_, *θ*_2*i*_), *i* = 1, …, *m* denote the joint hidden states across two studies. Then *τ*_*i*_ ∈ *{*(0, 0), (0, 1), (1, 0), (1, 1)*}* with ℙ (*τ*_*i*_ = (*k, l*)) = *ξ*_*kl*_ for *k, l* ∈ *{*0, 1*}*, and 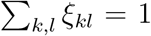. Here *ξ*_00_, *ξ*_01_, *ξ*_10_ and *ξ*_11_ denote the probabilities of hidden states (0, 0), (0, 1), (1, 0) and (1, 1), respectively. Based on this four-group model, the replicability null hypothesis for *i*th gene can be defined as

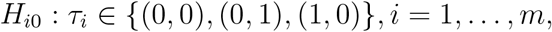

where the *i*th gene is replicable if it shows expression patterns in both study 1 and study 2. For simplicity, we denote the hidden state of *H*_*i*0_ as *h*_*i*_, where *h*_*i*_ = 0 indicates *H*_*i*0_ is true and *h*_*i*_ = 1 otherwise. Hence ℙ (*h*_*i*_ = 0) = *ξ*_00_ + *ξ*_01_ + *ξ*_10_ and ℙ(*h*_*i*_ = 1) = *ξ*_11_.

### 2.2 JUMP for replicability analysis

The schematic of JUMP for identifying replicable SVGs from two SRT studies is shown in Fig. 1. After obtaining paired *p*-values from two studies, we define the maximum *p*-values as

**Figure 1.**
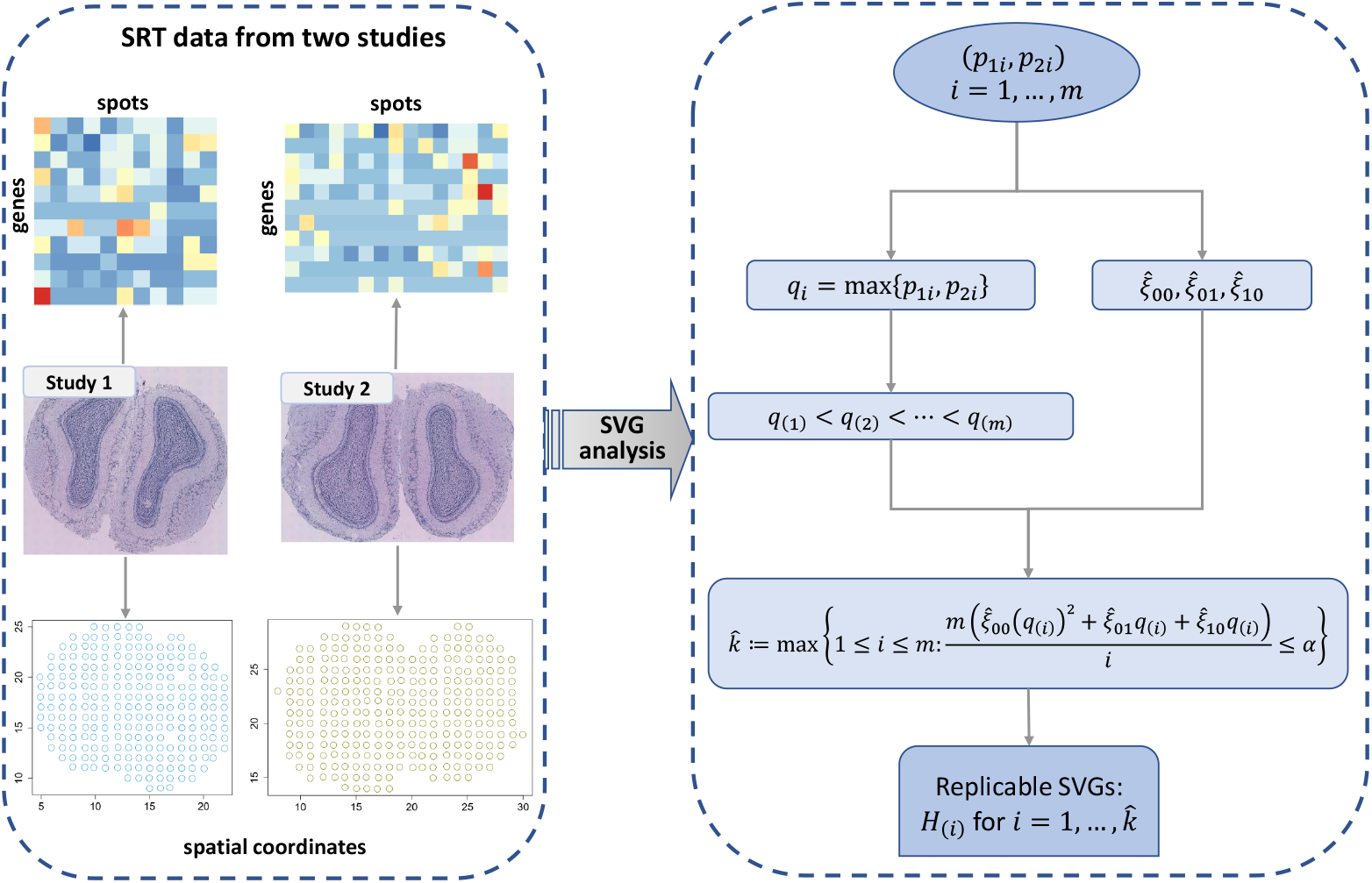
Schematic of JUMP for identifying replicable SVGs from two SRT studies

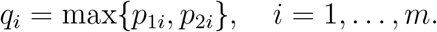

Under the assumption that *f*_0_ follows the standard uniform distribution, we have

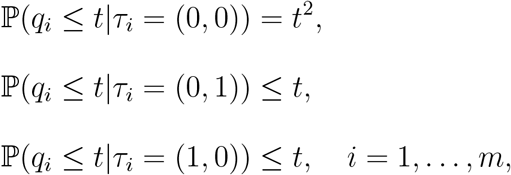

where we use the assumption that conditional on the joint hidden states, two *p*-value sequences are independent.

Under the replicable null, *q*_*i*_ follows a two-group mixture model

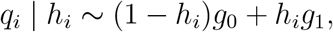

where *g*_0_ and *g*_1_ denote the density function of *q*_*i*_ under replicable null and non-null, respectively. For any *t* ∈ (0, 1), we compute the cumulative distribution function of *q*_*i*_ under replicable null as follows.

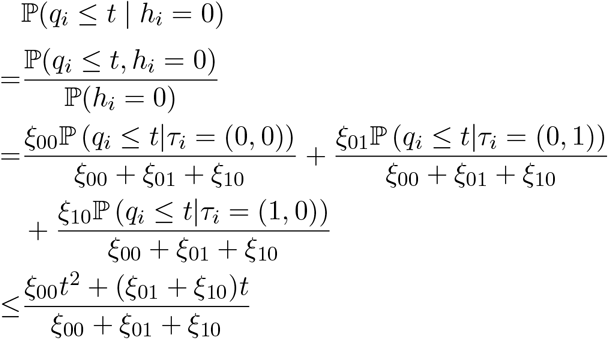

Denote

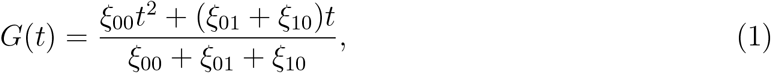

we have

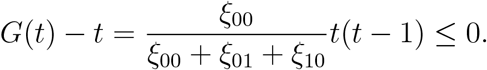

Hence ℙ (*q*_*i*_ *≤ t* | *h*_*i*_ = 0) *≤ t*, which means that *q*_*i*_ follows a super-uniform distribution under the replicability null. This verifies that the vanilla maximum *p*-value method has valid FDR control.

For *t* ∈ (0, 1), the number of rejections is 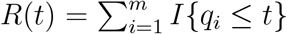, and the number of false rejections is bounded by *m*(*ξ*_00_ +*ξ*_01_ +*ξ*_10_)*G*(*t*). At threshold *t*, we have a conservative estimate of FDR by

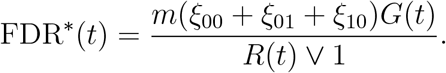

In the oracle case that we know *ξ*_00_, *ξ*_01_ and *ξ*_10_, at FDR level *α*, let

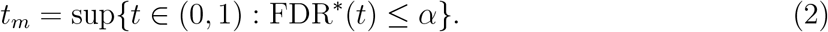

Reject *H*_*i*_ if *q*_*i*_ *≤ t*_*m*_.

### 2.3 Estimation of unknowns

As *ξ*_00_, *ξ*_01_, and *ξ*_10_ are unknown in practice, we provide their estimates in this section. Following Storey (2002) and Storey et al. (2004), in study *j*, the proportion of null hypotheses, 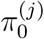, can be estimated by

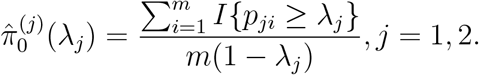

Similarly, we estimate *ξ*_00_ as

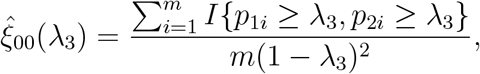

where *λ*_1_, *λ*_2_ and *λ*_3_ are tuning parameters. We use the smoothing method provided in Storey and Tibshirani (2003) to select tuning parameters. Please see Supplementary Note A for more details. We have

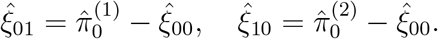

We estimate FDR^*∗*^(*t*) by

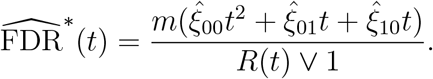

The data-adaptive thresholding is

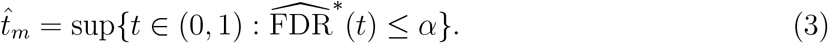

We claim the replicability of *i*th gene if 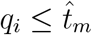.

This is equivalent to the following step-up procedure. First, order the maximum *p*-values *q*_(1)_ *≤ … ≤ q*_(*m*)_. Second, find

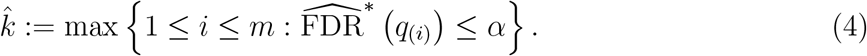

Reject *H*_(*i*)_ for 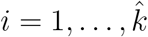, where *H*_(*i*)_ corresponds to *q*_(*i*)_.

The key to the power gain is to incorporate different states in the composite null. This is similar in spirit to plugging in the proportion of the null hypothesis with a single *p*-value sequence in Storey (2002).

## 3 Results

In this section, we evaluate the FDR control and power of JUMP via simulations and conduct data analysis to identify replicable SVGs from two pairs of SRT datasets measured with different technologies.

### 3.1 Simulation studies

We conducted extensive simulation studies to evaluate the performance of different methods. Power is defined as the expectation of true replicable discoveries over the total number of non-null hypotheses. We compare JUMP with *ad hoc* BH, MaxP, radjust (Bogomolov and Heller, 2018), MaRR (Philtron et al., 2018), and IDR (Li et al., 2011) for replicability analysis. In addition, we used two meta-analysis methods, Šidák’s method (Šidák, 1967) and Lancaster’s method (Lancaster, 1961), that combine *p*-values across two studies. We applied the BH procedure (Benjamini and Hochberg, 1995) on the aggregated *p*-values to show they are not suitable for replicability analysis. Detailed description of different methods can be found in Supplementary Note B.

In each simulation, states of genes in two studies, *θ*_1*i*_ and *θ*_2*i*_, were generated from a multinomial distribution with probabilities, ℙ(*θ*_1*i*_ = *k, θ*_2*i*_ = *l*) = *ξ*_*kl*_, *k, l* ∈ *{*0, 1*}*, for prespecified *ξ*_00_, *ξ*_01_, *ξ*_10_ and *ξ*_11_. Denote *N* (*µ, σ*^2^) as a normal distribution with mean *µ* and standard deviation *σ*. In simulation study 1, we independently generated summary statistic 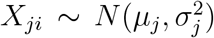 for *i*th gene in study *j* (*j* = 1, 2), where *µ*_*j*_ = 0 if *θ*_*ji*_ = 0, and *µ*_*j*_ *>* 0 if *θ*_*ji*_ = 1. One-sided *p*-values for study *j* were obtained by *p*_*ji*_ = 1 *−* Φ(*Z*_*ji*_), *i* = 1, …, *m*, where *Z*_*ji*_ = *X*_*ji*_*/σ*_*j*_ denotes the *Z*-statistic for the *i*th gene in study *j* and Φ(·) denotes the cumulative distribution function of *N* (0, 1). Fig. 2 presents the FDR control and power comparison of different methods under the setting of *m* = 10, 000, *ξ*_11_ = 0.05 and *ξ*_01_ = *ξ*_10_ over different values of *ξ*_00_, *µ*_*j*_ and *σ*_*j*_, *j* = 1, 2. For a given value of *ξ*_00_, corresponding *ξ*_01_ and *ξ*_10_ can be obtained by *ξ*_01_ = *ξ*_10_ = (1 *− ξ*_00_ *− ξ*_11_)*/*2. At a target FDR level of 0.05, we observe that the Šidák, the Lancaster, the IDR and the *ad hoc* BH do not have valid FDR control. The MaxP, radjust, MaRR and JUMP controlled the FDR at 0.05 across all settings. MaxP is overly conservative, radjust and MaRR have decent power, and JUMP has the highest power across all settings.

**Figure 2.**
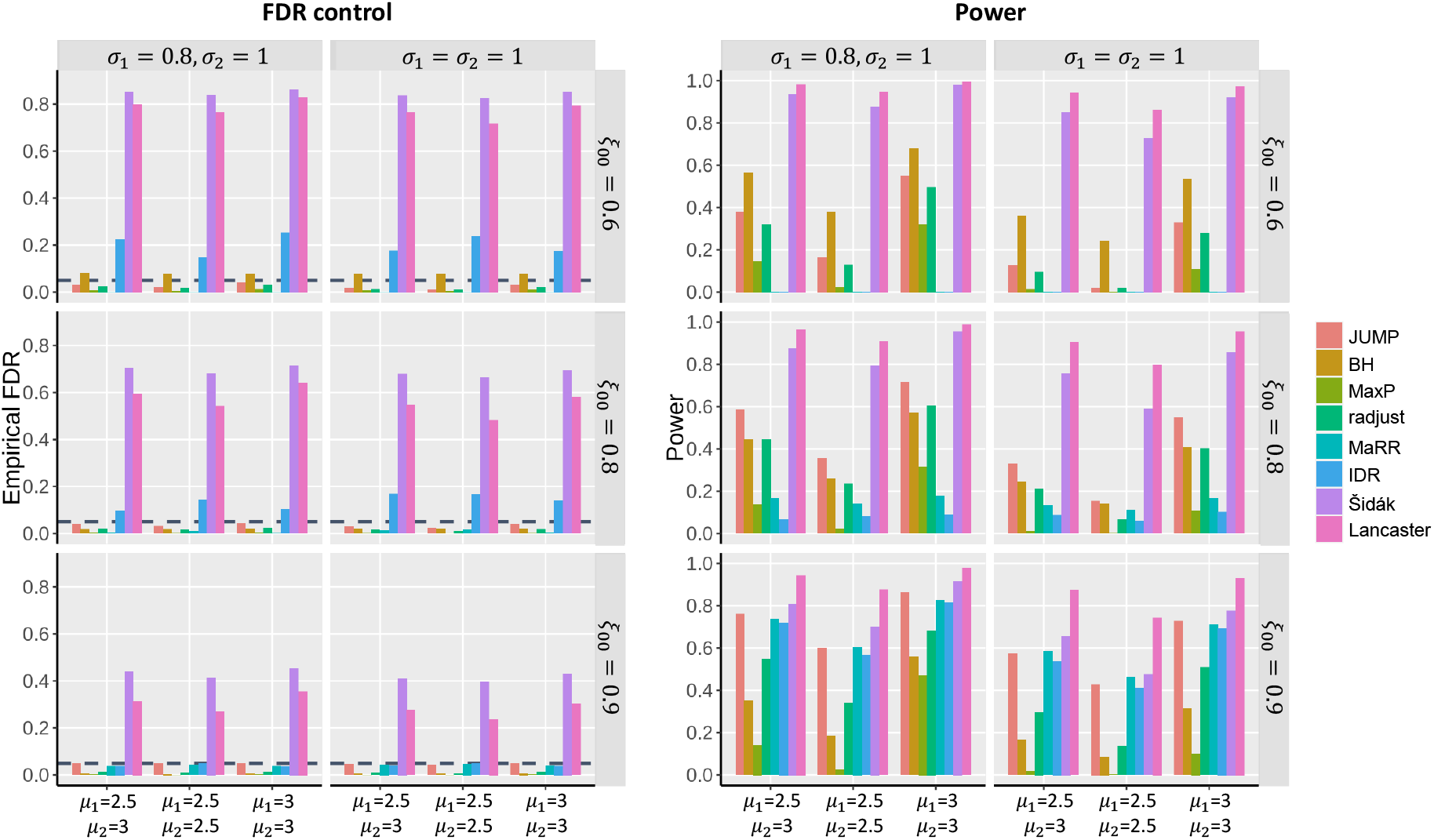
FDR control and power comparison of different methods in simulation studies. *m* = 10, 000, *ξ*_11_ = 0.05 and *ξ*_01_ = *ξ*_10_. Each row corresponds to different *ξ*_00_. Each column corresponds to different standard deviations. In each panel, the empirical FDR for different methods was calculated over 100 replications at a target FDR level of 0.05 (horizontal dashed line in the plots) for different non-null settings (left: *µ*_1_ = 2.5, *µ*_2_ = 3; middle: *µ*_1_ = *µ*_2_ = 2.5; right: *µ*_1_ = *µ*_2_ = 3).

We also performed realistic simulations based on Replicate 9 and Replicate 12 of ST datasets measured in mouse olfactory bulb (Ståhl et al., 2016). Details of the data generation process and simulation results can be found in Supplementary Note C, Fig. S1, Fig. S2 and Fig. S3.

### 3.2 Analysis of mouse olfactory bulb data

We first analyzed the SRT data from mouse olfactory bulb measured with ST technology (Ståhl et al., 2016) and 10X Visium technology. Ståhl et al. (2016) published 12 replicates of the mouse olfactory bulb ST datasets on the Spatial Research Website (https://www.spatialresearch.org/). We used Replicate 9 for our analysis, which includes 15, 284 genes on 237 spots. The 10X Visium dataset was downloaded from the 10X Visium spatial gene expression repository (https://www.10xgenomics.com/resources/datasets) and contains 32, 285 genes on 1, 185 spots. We filtered out genes that are expressed in less than 10% spatial spots and selected spots with at least ten total read counts, resulting in 9, 547 genes on 236 spots for the ST dataset and 10, 680 genes on 1, 185 spots for the 10X Visium dataset, respectively. We separately analyzed the two datasets using SPARK (Sun et al., 2020) to produce *p*-values. Paired *p*-values of 8, 547 genes common to both studies is the input to the replicability analysis. As can be seen in Fig. 3a, JUMP has higher power than the other two methods. At the FDR level of 0.05, MaxP identified 618 replicable SVGs, which were all detected by BH and JUMP. JUMP identified 807 replicable SVGs, including 637 of the 638 SVGs detected by BH. This is consistent with the simulation results that MaxP is overly conservative and JUMP has higher power.

**Figure 3.**
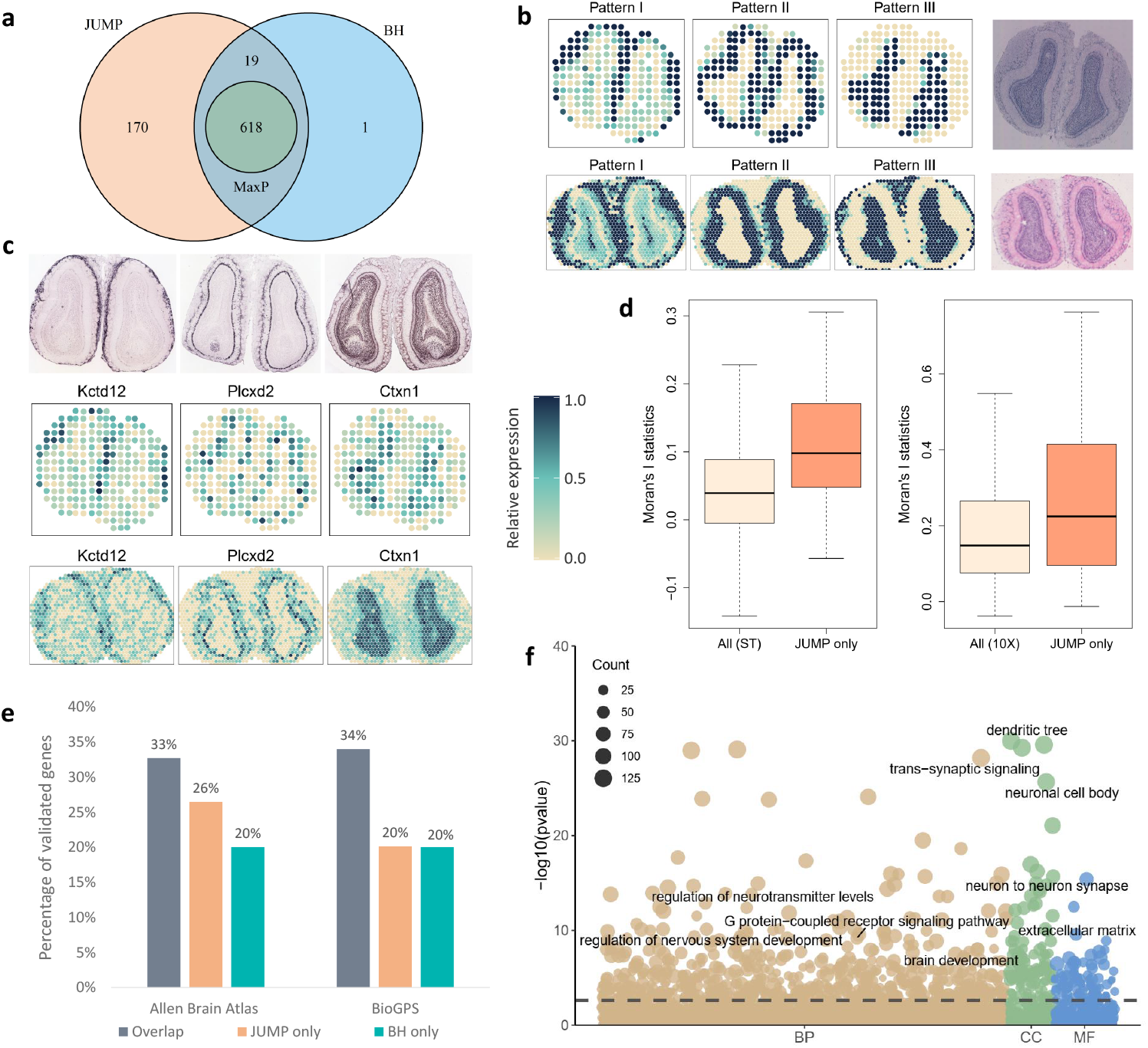
Analysis and validation results of the mouse olfactory bulb data measured with ST technology and 10X Visium technology. (a) Venn diagram shows the number of replicable SVGs identified by different methods at FDR level 0.05 and the intersection of discoveries. (b) Three distinct spatial expression patterns based on the 807 replicable SVGs identified by JUMP in the ST study (top) and 10X Visium study (bottom). Color represents relative expression levels (antique white: low; navy blue: high). The corresponding hematoxylin and eosin staining (HE) images for the two studies are shown in the right panel. (c) Spatial expression patterns of three representative genes identified by JUMP, corresponding to Pattern I-III, respectively. *In situ* hybridization images of corresponding genes obtained from the Allen Brain Atlas are shown in the top panel. (d) The box plot shows Moran’s *I* statistic of the 189 replicable SVGs additionally identified by JUMP and that of all genes based on the ST study (left) and the 10X Visium study (right). (e) The bar chart displays the number of replicable SVGs additionally identified by JUMP and BH compared to that identified by all three methods. They were validated in two reference gene lists from the Harmonizome database: one from the Allen Brain Atlas dataset and the other from the BioGPS dataset. (f) The bubble plot shows the GO enrichment analysis result of JUMP. The horizontal dashed line represents the FDR level 0.01. The color of a bubble shows different GO term categories: BP (tan), CC (green), and MF (blue). The size of a bubble represents the counts of corresponding gene sets.

We clustered the 807 replicable SVGs identified by JUMP into three groups using the hierarchical agglomerative clustering algorithm implemented in the R package *amap* (v0.8-18) and summarized the spatial expression patterns based on the expression level of the genes in corresponding groups. In both studies, the summarized patterns were consistent with three main cell layers in mouse olfactory bulb. In Fig. 3b and Supplementary Fig. S4a, Pattern I corresponds to the glomerular layer, Pattern II corresponds to the mitral layer and Pattern III corresponds to the granular layer. Spatial expression patterns of three representative SVGs (*Kctd12, Plcxd2, Ctxn1*) identified by JUMP are presented in Fig. 3c, which correspond to Pattern I-III, respectively and are consistent with the *in situ* hybridization images obtained from the Allen Brain Atlas. For comparison, we also randomly selected 30 genes from the 189 additional findings of JUMP compared to the overlaps of three methods and showed their spatial expression patterns in Supplementary Fig. S5a, b. Moreover, we calculated Moran’s *I* statistic (Moran, 1950) of the 189 replicable SVGs additionally identified by JUMP and compared it with that using all 8, 547 genes to show the autocorrelations of the additional findings by JUMP (Fig. 3d).

To further compare and validate the replicable SVGs identified by different methods, we downloaded two published gene sets from the Harmonizome database (Rouillard et al., 2016) consisting of genes related to mouse olfactory bulb as a reference (Fig. 3e). The first gene set was summarized based on three layers (glomerular, mitral, and granular) of the main olfactory bulb from the Allen Brain Atlas adult mouse brain tissue gene expression profiles dataset (Sunkin et al., 2012), including 3, 485 genes with high or low expression in main olfactory bulb relative to other tissues. 33% of the 618 replicable SVGs that were identified by all three methods were validated. Among the 189 replicable SVGs additionally identified by JUMP, 26% were validated in the reference list, whereas only 4 of the 20 SVGs additionally identified by BH were in the same list. The second reference gene set includes 2, 031 genes differentially expressed in mouse olfactory bulb relative to other cell types and tissues from the BioGPS mouse cell type and tissue gene expression profiles dataset (Wu et al., 2013). In addition to the replicable SVGs identified by all three methods (34% validated), 38 of the 189 replicable SVGs only detected by JUMP were in the list, whereas 4 of the 20 replicable SVGs only detected by BH were validated in the same list. These results provide additional biological evidence for the improved power of JUMP.

Additionally, we performed Gene Ontology (GO) enrichment analysis on replicable SVGs identified by JUMP and BH (Fig. 3f). At the FDR level of 0.01, JUMP enriched 846 GO terms and BH enriched 764 GO terms (708 overlaps). Many of the 138 GO terms only identified by JUMP are related to nervous system development and olfactory bulb organization. For instance, transmission of nerve impulse (GO:0019226), neuronal action potential (GO:0019228), and forebrain neuron differentiation (GO:0021879) are closely related to the establishment of axodendritic and dendrodendritic synaptic contacts within the olfactory bulb (Belluzzi et al., 2003); GABA-ergic synapse (GO:0098982) plays a key role in the organization of olfactory bulb (Hanson et al., 2020); GO terms GO:0045744 and GO:0002029 are related to G protein-coupled receptor, which can be encoded by odorant receptor genes differentially expressed at conserved positions in the olfactory bulb (Katidou et al., 2018).

### 3.3 Analysis of mouse cerebellum data

We next analyzed the SRT data from mouse cerebellum measured with Slide-seq technology (Rodriques et al., 2019) and Slide-seqV2 technology (Stickels et al., 2021). The two datasets were obtained from Broad Institute’s single-cell repository (https://singlecell.broadinstitute.org/single_cell) with IDs SCP354 and SCP948, respectively. The Slide-seq dataset (file “Puck_180430_6”) contains 18, 671 genes on 25, 551 beads. we filtered out beads with total counts less than 50. The Slide-seqV2 dataset contains 23, 096 genes on 39, 496 beads. We cropped regions of interest by filtering out beads with total counts less than 100. Mitochondrial genes and genes that are not expressed in any locations were filtered out from the two datasets, resulting in 17, 481 genes on 14, 667 beads for the Slide-seq data and 20, 117 genes on 11, 626 beads for the Slide-seqV2 data. After applying the SPARK-X method (Zhu et al., 2021) on the two datasets separately, we obtained two sequences of *p*-values and matched them by gene. We used the paired *p*-values for 16, 519 genes common to both studies as input for the replicability analysis of SVGs. At FDR level 0.05, MaxP identified 279 replicable SVGs, which were all identified by BH and JUMP. JUMP detected all BH findings (394) and identified 54 additional replicable SVGs.

We first examined the spatial expression patterns of the 448 replicable SVGs identified by JUMP by clustering them into three groups showing distinct spatial expression patterns (Fig. 4a). In both Slide-seq (left) and Slide-seqV2 (right) datasets, the 448 SVGs showed consistent patterns: Pattern I and Pattern III correspond to the purkinje cell layer and granular cell layer, respectively; and Pattern II corresponds to other cell layers. Three representative genes identified by JUMP (*Pcp2, Mbp*, and *Snap25*) exhibited corresponding spatial expression patterns, which were validated by *in situ* hybridization in the Allen Brain Atlas (Fig. 4b). Supplementary Fig. S6a,b display spatial expression patterns of 24 genes randomly selected from the 169 replicable SVGs identified by JUMP in addition to genes identified by all three methods. As shown in Fig. 4d, the additional spatial autocorrelations revealed by JUMP were further confirmed by Moran’s *I* (Moran, 1950).

**Figure 4.**
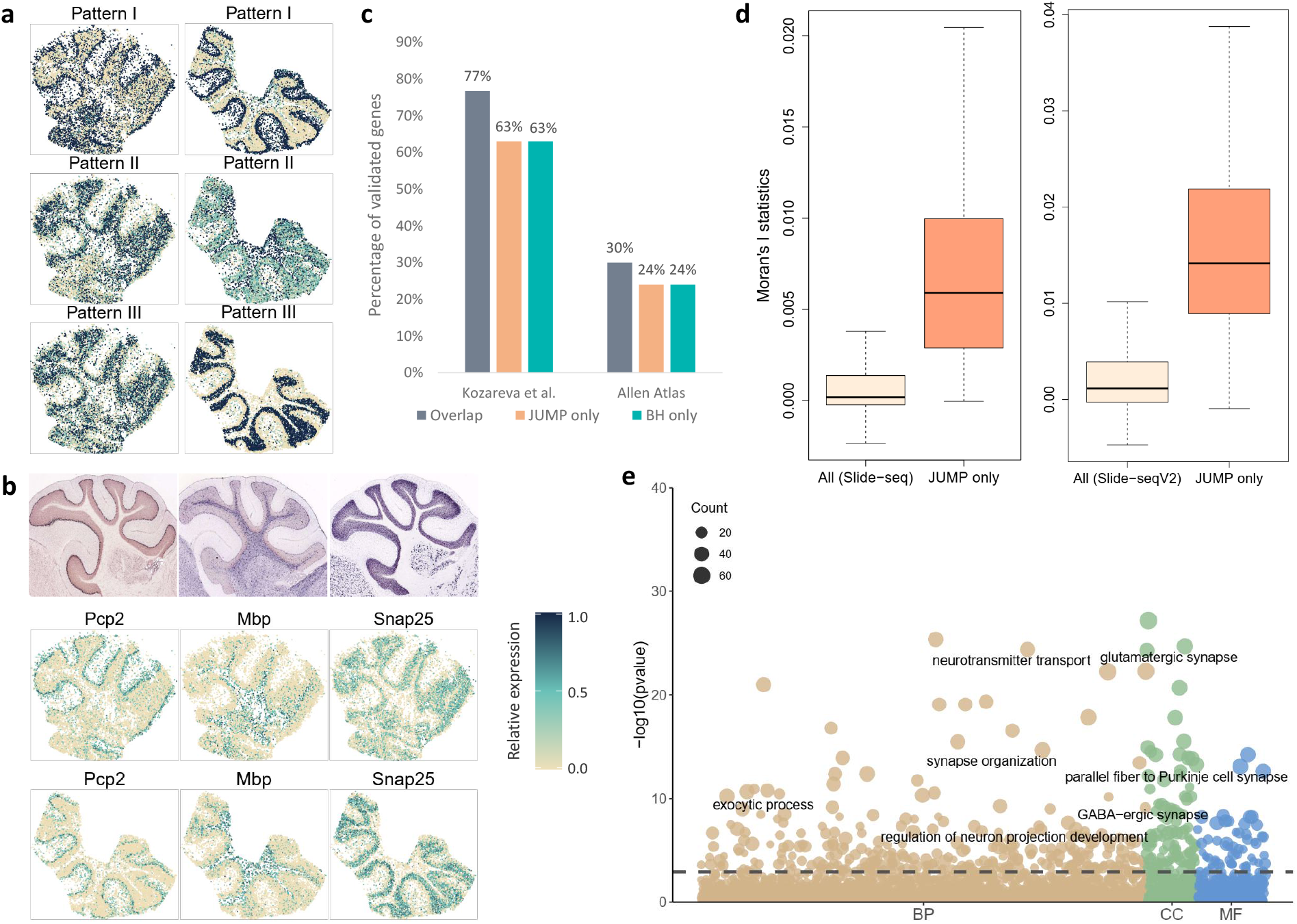
Analysis and validation results of the mouse cerebellum data measured with Slide-seq technology and Slide-seqV2 technology. (a) Three distinct spatial expression patterns based on the 448 replicable SVGs identified by JUMP in the Slide-seq study (left) and Slide-seqV2 study (right). Color represents relative expression levels (antique white: low; navy blue: high). (b) Spatial expression patterns of three representative genes identified by JUMP, corresponding to Pattern I-III, respectively. *In situ* hybridization images of corresponding genes obtained from the Allen Brain Atlas are shown in the top panel. (c) The bar chart displays the number of replicable SVGs additionally identified by JUMP and BH compared to that identified by all three methods. They were validated in two reference gene lists: one from Kozareva et al. (2021) and the other from the Allen Brain Atlas dataset summarized in the Harmonizome database. (d) The box plot shows Moran’s *I* statistic of the 169 replicable SVGs additionally identified by JUMP and that of all genes based on the Slide-seq study (left) and the Slide-seqV2 study (right). (e) The bubble plot shows the GO enrichment analysis result of JUMP. The horizontal dashed line represents the FDR level 0.01. The color of a bubble shows different GO term categories: BP (tan), CC (green), and MF (blue). The size of a bubble represents the counts of corresponding gene sets.

We provided two gene sets obtained from published literature to validate the results of different methods. First, we obtained a list of genes that are highly differentially expressed across all clusters in mouse cerebellar cortex from Kozareva et al. (2021). We further filtered out genes with absolute log fold change smaller than 0.05 and obtained 3, 976 genes for the validation. In addition to genes identified by all three methods (77% validated), 107 of the 169 replicable SVGs additionally identified by JUMP were in the list, whereas 73 of the 115 findings by BH were validated. Second, we downloaded three gene sets related to mouse cerebellum in the Allen Brain Atlas datasets (Sunkin et al., 2012) from the Harmonizome database (Rouillard et al., 2016) and summarized them to a list of 3, 000 genes that are differentially expressed in mouse cerebellum, cerebellar cortex, and cerebellar hemisphere. Among the 279 SVGs identified by all three methods, 30% were in this gene list. 39 of the 169 replicable SVGs additionally identified by JUMP were validated, whereas 28 of the 115 SVGs additionally identified by BH were in the same list.

Finally, we performed GO enrichment analysis on the replicable SVGs identified by different methods to evaluate the biological significance additionally identified by JUMP. At the FDR level of 0.01, JUMP enriched 452 GO terms and BH identified 418 GO terms (394 overlaps). Among the 58 GO terms only enriched by JUMP, many are closely related to the structural constitution and functional development of mouse cerebellum, such as GO terms of cerebellar cortex development (GO:0021695), cerebellum development (GO:0021549), central nervous system neuron development (GO0021954), dendritic transport (GO:0098935), and photoreceptor ribbon synapse (GO:0098684), among others.

## 4 Discussion

We present a new method, JUMP, for identifying replicable features from two high-throughput experiments. Analysis of different SRT studies identifies important replicable SVGs that might otherwise be missed by existing methods. JUMP is simple to implement and computationally scalable to tens of thousands of genes (Supplementary Table S1 and S2). Moreover, JUMP does not require two SRT studies to have the same spatial alignment or the same tissue thickness. In addition, JUMP is flexible and can accommodate data from other modalities, such as scRNA-seq, ATAC-seq, and CITE-seq, among others.

One limitation of JUMP is that it only identifies replicable features from two high-throughput experiments. If we want to extend it to more than two studies, say *n* studies, we require a 2^*n*^-group model for the data, and the composite null is composed of (2^*n*^ *−* 1) hidden joint states whose proportions need to be estimated. We leave this for future research.

## Data availability

The data underlying this article are publicly available in corresponding repositories. The mouse olfactory bulb ST dataset can be obtained with file ‘MOB Replicate 9’ in the Spatial Research Website at https://www.spatialresearch.org/resources-published-datasets/doi-10-1126science-aaf2403/. The mouse olfactory bulb 10X Visium dataset is available in the 10X Visium spatial gene expression repository at https://www.10xgenomics.com/resources/datasets/adult-mouse-olfactory-bulb-1-standard-1. The mouse cerebellum Slide-seq dataset and Slide-seqV2 dataset are provided in the Broad Institute’s single-cell repository at https://singlecell.broadinstitute.org/single_cell/ under IDs SCP354 (file ‘Puck_180430_6’) and SCP948, respectively.

## Acknowledgements

This work was partially supported by the China Postdoctoral Science Foundation (No. 801212021410) to Y. L.

## References

O. Belluzzi, M. Benedusi, J. Ackman, and J. J. LoTurco. Electrophysiological differentiation of new neurons in the olfactory bulb. Journal of Neuroscience, 23(32):10411–10418, 2003.

Y. Benjamini and Y. Hochberg. Controlling the false discovery rate: a practical and powerful approach to multiple testing. Journal of the Royal Statistical Society: Series B (Methodological), 57(1):289–300, 1995.

Y. Benjamini and D. Yekutieli. The control of the false discovery rate in multiple testing under dependency. Annals of Statistics, 29(4):1165–1188, 2001.

Y. Benjamini, R. Heller, and D. Yekutieli. Selective inference in complex research. Philosophical Transactions of the Royal Society A: Mathematical, Physical and Engineering Sciences, 367(1906):4255–4271, 2009.

M. Bogomolov and R. Heller. Discovering findings that replicate from a primary study of high dimension to a follow-up study. Journal of the American Statistical Association, 108(504): 1480–1492, 2013.

M. Bogomolov and R. Heller. Assessing replicability of findings across two studies of multiple features. Biometrika, 105(3):505–516, 2018.

M. Bogomolov and R. Heller. Replicability across multiple studies. arXiv preprint arXiv:2210.00522, 2022.

D. Chung, C. Yang, C. Li, J. Gelernter, and H. Zhao. Gpa: a statistical approach to prioritizing gwas results by integrating pleiotropy and annotation. PLoS Genetics, 10(11):e1004787, 2014.

D. Edsgärd, P. Johnsson, and R. Sandberg. Identification of spatial expression trends in single-cell gene expression data. Nature Methods, 15(5):339–342, 2018.

R. Fisher. Statistical Methods for Research Workers. Edinburgh Oliver & Boyd, 1925.

E. Hanson, J. Swanson, and B. R. Arenkiel. Gabaergic input from the basal forebrain promotes the survival of adult-born neurons in the mouse olfactory bulb. Frontiers in Neural Circuits, 14:17, 2020.

J. Hu, X. Li, K. Coleman, A. Schroeder, N. Ma, D. J. Irwin, E. B. Lee, R. T. Shinohara, and M. Li. Spagcn: Integrating gene expression, spatial location and histology to identify spatial domains and spatially variable genes by graph convolutional network. Nature Methods, 18 (11):1342–1351, 2021.

K. Hung and W. Fithian. Statistical methods for replicability assessment. The Annals of Applied Statistics, 14(3):1063–1087, 2020.

M. Katidou, X. Grosmaitre, J. Lin, and P. Mombaerts. G-protein coupled receptors mc4r and drd1a can serve as surrogate odorant receptors in mouse olfactory sensory neurons. Molecular and Cellular Neuroscience, 88:138–147, 2018.

I. Kleino, P. Frolovaitė, T. Suomi, and L. L. Elo. Computational solutions for spatial transcriptomics. Computational and Structural Biotechnology Journal, 2022.

V. Kozareva, C. Martin, T. Osorno, S. Rudolph, C. Guo, C. Vanderburg, N. Nadaf, A. Regev, W. G. Regehr, and E. Macosko. A transcriptomic atlas of mouse cerebellar cortex comprehensively defines cell types. Nature, 598(7879):214–219, 2021.

H. Lancaster. The combination of probabilities: an application of orthonormal functions. Australian Journal of Statistics, 3(1):20–33, 1961.

Q. Li, J. B. Brown, H. Huang, and P. J. Bickel. Measuring reproducibility of high-throughput experiments. Annals of Applied Statistics, 5(3):1752–1779, 2011.

P. A. Moran. Notes on continuous stochastic phenomena. Biometrika, 37(1/2):17–23, 1950.

D. Philtron, Y. Lyu, Q. Li, and D. Ghosh. Maximum rank reproducibility: a nonparametric approach to assessing reproducibility in replicate experiments. Journal of the American Statistical Association, 113(523):1028–1039, 2018.

S. G. Rodriques, R. R. Stickels, A. Goeva, C. A. Martin, E. Murray, C. R. Vanderburg, J. Welch, L. M. Chen, F. Chen, and E. Z. Macosko. Slide-seq: A scalable technology for measuring genome-wide expression at high spatial resolution. Science, 363(6434):1463–1467, 2019.

A. D. Rouillard, G. W. Gundersen, N. F. Fernandez, Z. Wang, C. D. Monteiro, M. G. Mc-Dermott, and A. Ma’ayan. The harmonizome: a collection of processed datasets gathered to serve and mine knowledge about genes and proteins. Database, 2016.

Z. Šidák. Rectangular confidence regions for the means of multivariate normal distributions. Journal of the American Statistical Association, 62(318):626–633, 1967.

P. L. Ståhl, F. Salmén, S. Vickovic, A. Lundmark, J. F. Navarro, J. Magnusson, S. Giacomello, M. Asp, J. O. Westholm, M. Huss, et al. Visualization and analysis of gene expression in tissue sections by spatial transcriptomics. Science, 353(6294):78–82, 2016.

R. R. Stickels, E. Murray, P. Kumar, J. Li, J. L. Marshall, D. J. Di Bella, P. Arlotta, E. Z. Macosko, and F. Chen. Highly sensitive spatial transcriptomics at near-cellular resolution with slide-seqv2. Nature Biotechnology, 39(3):313–319, 2021.

J. D. Storey. A direct approach to false discovery rates. Journal of the Royal Statistical Society: Series B (Statistical Methodology), 64(3):479–498, 2002.

J. D. Storey and R. Tibshirani. Statistical significance for genomewide studies. Proceedings of the National Academy of Sciences, 100(16):9440–9445, 2003.

J. D. Storey, J. E. Taylor, and D. Siegmund. Strong control, conservative point estimation and simultaneous conservative consistency of false discovery rates: a unified approach. Journal of the Royal Statistical Society: Series B (Statistical Methodology), 66(1):187–205, 2004.

S. Sun, J. Zhu, and X. Zhou. Statistical analysis of spatial expression patterns for spatially resolved transcriptomic studies. Nature Methods, 17(2):193–200, 2020.

S. M. Sunkin, L. Ng, C. Lau, T. Dolbeare, T. L. Gilbert, C. L. Thompson, M. Hawrylycz, and C. Dang. Allen brain atlas: an integrated spatio-temporal portal for exploring the central nervous system. Nucleic Acids Research, 41(D1):D996–D1008, 2012.

V. Svensson, S. A. Teichmann, and O. Stegle. Spatialde: identification of spatially variable genes. Nature Methods, 15(5):343–346, 2018.

C. Wu, I. MacLeod, and A. I. Su. Biogps and mygene. info: organizing online, gene-centric information. Nucleic Acids Research, 41(D1):D561–D565, 2013.

J. Zhu, S. Sun, and X. Zhou. Spark-x: non-parametric modeling enables scalable and robust detection of spatial expression patterns for large spatial transcriptomic studies. Genome Biology, 22(1):1–25, 2021.

