## Supplementary for "JUMP: replicability analysis of high-throughput experiments with applications to spatial transcriptomic studies"

### Supplementary Materials for "JUMP: replicability analysis of high-throughput experiments with applications to spatial transcriptomic studies"

Pengfei Lyu[1†], Yan Li[2†], Xiaoquan Wen[3] and Hongyuan Cao[1*]

[1]Department of Statistics, Florida State University, Tallahassee, FL 32306, USA. [2]School of Mathematics, Jilin University, Changchun, Jilin 130012, China. [3]Department of Biostatistics, University of Michigan, Ann Arbor, MI 48109, USA.

[†]The first two authors should be regarded as Joint First Authors.

### A General algorithm for estimating the unknowns

JUMP requires tuning parameters $\lambda_1, \lambda_2$ and $\lambda_3$ in estimation of the unknown parameters, $\hat{\pi}_0^{(1)}(\lambda_1), \hat{\pi}_0^{(2)}(\lambda_2)$ and $\hat{\xi}_{00}(\lambda_3)$, which involves a trade-off between bias and variance. We use the following general algorithm provided in [9] to estimate $\hat{\pi}_0^{(1)}(\lambda_1), \hat{\pi}_0^{(2)}(\lambda_2)$ and $\hat{\xi}_{00}(\lambda_3)$ from $p$-values, $(p_{1i}, p_{2i}), i = 1, \ldots, m$, respectively.

1. For a range of $\lambda_1, \lambda_2$, and $\lambda_3$, say $\{0.01, 0.02, 0.03, \ldots, 0.80\}$, calculate corresponding estimates by

$$\hat{\pi}_0^{(1)}(\lambda_1) = \frac{\sum_{i=1}^m I\{p_{1i} \geq \lambda_1\}}{m(1-\lambda_1)},$$
$$\hat{\pi}_0^{(2)}(\lambda_2) = \frac{\sum_{i=1}^m I\{p_{2i} \geq \lambda_2\}}{m(1-\lambda_2)},$$
$$\hat{\xi}_{00}(\lambda_3) = \frac{\sum_{i=1}^m I\{p_{1i} \geq \lambda_3, p_{2i} \geq \lambda_3\}}{m(1-\lambda_3)^2}.$$

2. Fit three natural cubic splines with 3 degrees of freedom for $\hat{\pi}_0^{(1)}(\lambda_1), \hat{\pi}_0^{(2)}(\lambda_2)$ and $\hat{\xi}_{00}(\lambda_3)$ over $\lambda_1, \lambda_2$ and $\lambda_3$, denoted as $\hat{f}_1, \hat{f}_2$ and $\hat{f}_3$, respectively.

3. Find $\hat{\lambda}_1, \hat{\lambda}_2$ and $\hat{\lambda}_3$ corresponding to zero derivatives of $\hat{f}_1, \hat{f}_2$ and $\hat{f}_3$. Let $\hat{\pi}_0^{(1)}(\hat{\lambda}_1), \hat{\pi}_0^{(2)}(\hat{\lambda}_2)$ and $\hat{\xi}_{00}(\hat{\lambda}_3)$ be our final estimates.

### B Comparison methods overview

In the simulation study, We compared JUMP to several statistical methods for replicability analysis methods (*ad hoc* BH, naïve MaxP, IDR, MaRR and radjust) and two $p$-value combination methods for meta-analysis. Let $(p_{1i}, p_{2i}), i = 1, \ldots, m$ denote the paired $p$-values from two studies. We review these comparison methods as follows.

#### B.1 The *Ad hoc* BH method

BH [1] is the most popular multiple testing procedure that conservatively controls the FDR for $m$ independent or positively correlated tests. In study $j, j = 1, 2$, the BH procedure proceeds as below:

- *Step 1.* Let $p_{j(1)} \leq p_{j(2)} \leq \cdots \leq p_{j(m)}$ be the ordered $p$-values in study $j$, and denote by $H_{(i)}^{(j)}$ the null hypothesis corresponding to $p_{j(i)}$;
- *Step 2.* Find the largest $i$ such that $p_{j(i)} \leq \frac{i}{m}\alpha$, i.e., $\hat{k} = \max\{i \geq 1 : p_{j(i)} \leq \frac{i}{m}\alpha\}$, and $\hat{k} = 0$ if the set is empty;
- *Step 3.* Reject all $H_{(i)}^{(j)}$ for $i = 1, \ldots, \hat{k}$.

The *ad hoc* BH method for replicability analysis identifies features rejected by both studies as replicable signals.

#### B.2 The naïve MaxP method

Define the maximum $p$-values as

$$q_i = \max\{p_{1i}, p_{2i}\}, i = 1, \ldots, m.$$

As discussed in the paper, $q_i$ follows a super-uniform distribution under the replicability null. The naïve MaxP method directly applies BH [1] to $q_i, i = 1, \ldots, m$ for FDR control of replicability analysis.

#### B.3 The radjust procedure

The radjust procedure [2] works as follows,

- *Step 1.* For a pre-specified FDR level $\alpha$, compute

$$R = \max \left[ r : \sum_{i \in \mathcal{S}_1 \cap \mathcal{S}_2} I \left\{ (p_{1i}, p_{2i}) \leq \left( \frac{r\alpha}{2|\mathcal{S}_2|}, \frac{r\alpha}{2|\mathcal{S}_1|} \right) \right\} = r \right],$$

  where $\mathcal{S}_j$ is the set of features pre-selected in study $j$ for $j = 1, 2$. By default, it selects features with $p$-values less than or equal to $\alpha/2$.

- *Step 2.* Declare as replicated the features with indices in the set

$$\mathcal{R} = \left\{ i : (p_{1i}, p_{2i}) \leq \left( \frac{R\alpha}{2|\mathcal{S}_2|}, \frac{R\alpha}{2|S_1|} \right), i \in \mathcal{S}_1 \cap \mathcal{S}_2 \right\}.$$

This procedure gains power by pre-filtering irrelevant features. It looks very similar to *ad-hoc* BH procedure where the BH procedure is implemented for each study and the intersection of significant findings are regarded as replicable features. After close inspection, we find these two procedures are quite different. We use the following toy example to illustrate the difference between *ad hoc* BH and radjust procedure.

Assume we have two lists of $p$-values

$$(p_{1i})_{i=1}^{10} = (0, 0, 0, 0, \alpha/12, 3\alpha/10, 1, 1, 1, 1),$$
$$(p_{2i})_{i=1}^{10} = (1, 1, 1, 1, 3\alpha/10, \alpha/12, 0, 0, 0, 0).$$

Applying BH procedure separately to the two studies with FDR level $\alpha/2$, we get the rejections $\mathcal{R}_1 = \{1, 2, 3, 4, 5, 6\}$ and $\mathcal{R}_2 = \{5, 6, 7, 8, 9, 10\}$. The discovery set of *ad hoc* BH is $\mathcal{R}_1 \cap \mathcal{R}_2 = \{5, 6\}$.

If we pre-select $p$-values that are less than or equal to $\alpha/2$, we get the pre-selection sets $\mathcal{S}_1 = \{1, 2, 3, 4, 5, 6\}$ and $\mathcal{S}_2 = \{5, 6, 7, 8, 9, 10\}$. At FDR level $\alpha/2$, implementing Step 1 and 2 of radjust, we obtain $R = 0$ and $\mathcal{R} = \emptyset$. This toy example shows that radjust is more conservative than the *ad hoc* BH procedure.

#### B.4 The IDR procedure

The IDR procedure [5] deals with high throughput experimental data from two studies. For feature $i$, we have the bivariate observations $(x_{1i}, x_{2i}), i = 1, \ldots, m$. It is assumed that $(x_{1i}, x_{2i}), i = 1, \ldots, m$ consist of genuine signals (replicable signals across two studies) and spurious signals (non-replicable signals). Let $K_i$ denote whether the $i$th feature is a replicable signal ($K_i = 1$) or not ($K_i = 0$). It is assumed that $K_i, i = 1, \ldots, m$ are independent and follow the Bernoulli distribution. Denote $\pi_1 = P(K_i = 1)$. To induce dependence between $x_{1i}$ and $x_{2i}$, we use a copula model. Specifically, we assume that the observed data $(x_{1i}, x_{2i})$ are generated from latent variables $(z_{1i}, z_{2i})$. The latent variables

$$\begin{pmatrix} z_{1i} \\ z_{2i} \end{pmatrix} \Bigg| K_i = k \sim N\left( \begin{pmatrix} \mu_k \\ \mu_k \end{pmatrix}, \sigma_k^2 \begin{pmatrix} 1 & \rho_k \\ \rho_k & 1 \end{pmatrix} \right), \quad k = 0, 1,$$

where $\mu_0 = 0, \mu_1 > 0, \sigma_0^2 = 1, \sigma_1^2 > 0, \rho_0 = 0$, and $0 < \rho_1 \leq 1$. The cdf of $z_{ji}$ is

$$G(x) = P(z_{ji} \leq x) = \pi_1 \Phi\left(\frac{x - \mu_1}{\sigma_1}\right) + (1 - \pi_1)\Phi(x).$$

Denote the marginal distribution function of $x_{ji}, i = 1, \ldots, m; j = 1, 2$, as $F_j$. Generate

$$x_{ji} = F_j^{-1}(G(z_{ji})), i = 1, \ldots, m; j = 1, 2.$$

In this way, dependence across two studies is produced. To control the false discovery rate, we use the local irreproducible discovery rate (idr) as the test statistic, which is defined as the posterior probability of $K_i = 0$ given $(x_{1i}, x_{2i})$. Specifically,

$$idr(x_{1i}, x_{2i}) := P(K_i = 0 \mid x_{1i}, x_{2i}) \\ = \frac{(1-\pi_1)h_0[G^{-1}\{F_1(x_{1i})\}, G^{-1}\{F_2(x_{2i})\}]}{(1-\pi_1)h_0[G^{-1}\{F_1(x_{1i})\}, G^{-1}\{F_2(x_{2i})\}] + \pi_1 h_1[G^{-1}\{F_1(x_{1i})\}, G^{-1}\{F_2(x_{2i})\}]}.$$

where

$$h_k \sim N\left(\begin{pmatrix}\mu_k \\ \mu_k\end{pmatrix}, \sigma_k^2 \begin{pmatrix}1 & \rho_k \\ \rho_k & 1\end{pmatrix}\right), \; k = 0, 1.$$

The estimation of $(\pi_1, \mu_1, \sigma_1^2, \rho_1)$ and $(F_1, F_2)$ is through the EM algorithm [3]. The step-up procedure based on ordered idr can be used for FDR control [11]. Specifically, let $idr_{(1)} \leq \cdots \leq idr_{(m)}$ be the ranked $idr$ values, and denote $H_{(1)}, \ldots, H_{(m)}$ as the corresponding hypotheses. Find $l = \max\{i : i^{-1} \sum_{j=1}^{i} idr_j \leq \alpha\}$, and reject all $H_{(i)}$ with $i = 1, \ldots, l$.

#### B.5 The MaRR procedure

The MaRR procedure [6] uses the maximum rank of each feature. The null hypothesis is that $H_{0i}$ : $p_{1i}$ and $p_{2i}$ are irreproducible. Denote $(R_{1i}, R_{2i})$ as the ranks of $(p_{1i}, p_{2i}), i = 1, \ldots, m$ within each study. Define

$$M_i = \max\{R_{1i}, R_{2i}\}, i = 1, \ldots, m.$$

Let $\pi_1$ denote the proportion of replicable signals. Under the assumptions:

(I1) if gene $g$ is reproducible and gene $h$ is irreproducible

$$R_{1g} < R_{1h}, \quad R_{2g} < R_{2h};$$

(I2) the correlation between the ranks of the reproducible gene is non-negative;

(I3) the two ranks of the irreproducible gene are independent,

irreproducible ranks $R_{1i}$ and $R_{2i}$ are uniformly distributed between $\lfloor m\pi_1 \rfloor + 1$ and $m$. Denote the conditional null survival function of $M_i/m$ as

$$
\begin{aligned}
S_{m,\pi_1}(x) &= P(M_i/m > x \mid \text{ gene } i \text{ is irreproducible}) \\
=&1 - P\left(R_{1i}/m \leq x, R_{2i}/m \leq x \mid \text{ gene } i \text{ is irreproducible}\right) \\
=&1 - \prod_{j=1}^{2} P\left(R_{ji}/m \leq x \mid \text{ gene } i \text{ is irreproducible}\right) \\
=&\begin{cases} 1, & x < \pi_1, \\ 1 - \frac{(i_x - j_{\pi_1})^2}{(m - j_{\pi_1})^2}, & \pi_1 \leq x \leq 1, \end{cases}
\end{aligned}
$$

where $i_x = \lfloor mx \rfloor$ and $j_{\pi_1} = \lfloor m\pi_1 \rfloor$. The limiting conditional survival function of $M_i/m$ under the null is

$$
S_{m,\pi_1}(x) \to S_{\pi_1}(x) = \begin{cases} 1 & x < \pi_1 \\ 1 - \frac{(x-\pi_1)^2}{(1-\pi_1)^2} & \pi_1 \leq x \leq 1 \\ 0 & 1 < x \end{cases} \cdot
$$

The empirical survival function can be estimated by $\hat{S}_m(x) = \frac{1}{m}\sum_{i=1}^{m} I(M_i/m \geq x)$, $x \in (0,1)$. By strong law of large numbers and Bayesian formula,

$$
\begin{aligned}
\hat{S}_m(x) \to& P(M_i/m \geq x) \\
&= (1-\pi_1)P(M_i/m \geq x \mid \text{ gene } i \text{ is irreproducible}) + \pi_1 \times 0 \\
&= (1-\pi_1)S_{\pi_1}(x) \text{ for } x \in (\pi_1, 1).
\end{aligned}
$$

If we estimate $\pi_1$ by $i/m$, we can define the mean square error (MSE) as follows.

$$\text{MSE}(i/m) = (m-i)^{-1} \sum_{j=i}^{m} \left( \hat{S}_m(j/m) - (1-i/m) S_{i/m}(j/m) \right)^2 .$$

$\hat{k}$ is chosen to minimize the MSE in the range between 0 and $\lfloor 0.9m \rfloor$.

$$\hat{k} = \underset{i=0,1,\ldots,\lfloor 0.9m \rfloor}{\arg\min} \left\{ \text{MSE}(i/m) \right\} .$$

Thus $\hat{k}/m$ serves as a good estimation of $\pi_1$. To control the FDR at level $\alpha$, the MaRR generates the rejection threshold as follows

$$\text{Define } \hat{N} = \max_{\hat{k} < i \leq n} \left\{ i : m\widehat{\text{FDR}}(i) = \frac{(i-\hat{k})^2}{Q(i)(m-\hat{k})} \leq \alpha \right\},$$

where $Q(i) = \sum_{j=1}^{m} I(M_j \leq i)$. Reject all features associated with $M_i \leq \hat{N}$. Philtron et al [6] relax assumption (I1) to (R1): $P(R_{1g} < R_{1h}) > 1/2$ and $P(R_{2g} < R_{2h}) > 1/2$, which is more plausible in practice.

#### B.6 The Šidák's method

The Šidák-corrected minimum $p$-value [7] can be used for meta-analysis. Specifically, we calculate the aggregated $p$-values across two studies through

$$q_i^S = 1 - (1 - \min\{p_{1i}, p_{2i}\})^2, i = 1, \ldots, m.$$

Assume that $p_{1i}$ and $p_{2i}, i = 1, \ldots, m$ are independent. Under the null for meta-analysis where $p_{1i}$ and $p_{2i}$ follow standard uniform distribution, we compute the cdf of $\min\{p_{1i}, p_{2i}\}$. Specifically, we have $P(\min\{p_{1i}, p_{2i}\} \leq t) = 1 - (1-t)^2$. Denote $F(t) = 1 - (1-t)^2$,

$q_i^S = F(\min\{p_{1i}, p_{2i}\})$ follows a standard uniform distribution under the meta-analysis null. Here we use the property that for a standard uniformly distributed random variable $U$, the cdf of $F^{-1}(U)$ is $F$.

We apply the BH procedure [1] on $q_i^S, i = 1, \ldots, m$ to evaluate the performance of Šidák's method in replicability analysis.

#### B.7 The Lancaster's method

Lancaster's method [4] uses different weights for different studies. Denote $F_{\chi^2_{w_j}}$ as the cdf of a $\chi^2$ distribution with $w_j, j = 1, 2$ degree of freedom. For the $i$th hypothesis, Lancaster's method combines information across two studies by a test statistic $L_i = \sum_{j=1}^{2} F^{-1}_{\chi^2_{w_j}}(p_{ji})$, which follows a $\chi^2$ distribution with degree of freedom $w_1 + w_2$ under the null for meta-analysis that both studies are from the null. The $p$-value for Lancaster's method is computed as the tail probability of the $\chi^2$ distribution with $w_1 + w_2$ degrees of freedom evaluated at $L_i$. We denote them as $q_i^L, i = 1, \ldots, m$.

We apply the BH procedure [1] on $q_i^L, i = 1, \ldots, m$ to evaluate the performance of Lancaster's method in replicability analysis.

### C Realistic simulation studies

We performed realistic simulations based on Replicate 9 and Replicate 12 of the mouse olfactory bulb data measured with ST technology (files 'MOB Replicate 9' and 'MOB Replicate 12' in the Spatial Research Website at `https://www.spatialresearch.org`) [8]. The two datasets include $15,284$ genes measured on 237 spatial spots and $16,034$ genes measured on 282 spots, respectively. We filtered out genes that are expressed in less than 10% spatial spots and selected spots with at least ten total read counts, resulting in $9,547$ genes on 236 spots for the Replicate 9 dataset and $9,904$ genes on 279 spots for the Replicate 12 dataset.

The spatial expression patterns and parameters used in data generation for each study were inferred from SPARK [10]. We separately generated SRT count data based on the two studies following the simulation design in [10].

In study $j$ ($j = 1, 2$), for each gene, the count on spot $i$ was generated from

$$
\begin{aligned}
y_i &\sim \text{Poisson}(N_i \lambda_i), \\
\log \lambda_i &= \beta_i + \epsilon_i,
\end{aligned}
\tag{S1}
$$

where $i = 1, \dots, 236$ for study 1 and $i = 1, \dots, 279$ for study 2; $N_i$ denotes total counts of all genes on spot $i$, which is obtained from the mouse olfactory bulb data [8]; $\lambda_i$ represents the relative expression level of the focused gene, which will be generated; $\beta_i$ is the mean value of $\log \lambda_i$; and $\epsilon_i \sim N(0, s_j^2)$ is the random noise. If the gene in focus is not spatially variable, we set $\beta_i$ across all spatial spots to be constant, which is the median value of intercepts estimated from SPARK [10]($-9.94$ for study 1 and $-9.93$ for study 2). If the focused gene is an SVG, we used different $\beta_i$ for spots to exhibit spatial expression patterns. Specifically, we first categorized the spots into two groups based on the three spatial expression patterns in Fig. S1: a group of spots with low expression levels and a group of spots with high expression levels. In the low expression group, we set $\beta_i$ to be the median value of intercepts estimated by SPARK ($-9.94$ for study 1 and $-9.93$ for study 2); in the high expression group, we set $\beta_i$ to be two-fold (weak signal strength), three-fold (moderate signal strength) or four-fold (strong signal strength) of the corresponding median value on rate parameter, e.g., $e^{\beta_i} = 2 \cdot e^a$ means $\beta_i$ is two-fold of $a$. Finally, $y_i$ was generated from (S1) with simulated $\beta_i$ and $\epsilon_i$.

Let $m = 10,000, \xi_{11} = 0.05$ and $\xi_{01} = \xi_{10}$. For a given value of $\xi_{00}$, corresponding $\xi_{01}$ and $\xi_{10}$ can be obtained by $\xi_{01} = \xi_{10} = (1 - \xi_{00} - \xi_{11})/2$. States of genes in two SRT studies, $\theta_{1i}$ and $\theta_{2i}$, were generated from a multinomial distribution with probabilities, $\mathbb{P}(\theta_{1i} = k, \theta_{2i} = l) = \xi_{kl}, k, l \in \{0, 1\}$, for pre-specified $\xi_{00}, \xi_{01}, \xi_{10}$ and $\xi_{11}$. After obtaining $\theta_{ji}$ for $i = 1, \dots, m$ and

$j = 1, 2$, we simulated gene count matrices based on corresponding ST data and parameters with different signal strengths (moderate or strong) and different standard deviations for the error term ($s_j = 0.3$ or $0.5$). Then we applied SPARK [10] on the two count data to get two paired $p$-values sequences, denoted as $(p_{1i}, p_{2i}), i = 1, \ldots, m$. Methods for replicability analysis are based on the paired $p$-value sequence.

Fig. S2 and Fig. S3 show the FDR control and power comparison of different methods across different settings. We observe that MaxP and JUMP controlled the FDR at the nominal level across all settings, and JUMP is more powerful than MaxP. BH is not valid in practice since it failed to control the FDR in some settings (e.g., $\xi_{00} = 0.5$). The power increased for all methods from Pattern I to Pattern III. By examining the three spatial expression patterns on which the corresponding data were generated (Supplementary Fig. S1), we speculate that this might be due to the increased spatial variability from Pattern I to Pattern III.

### D Computational time

We implemented all methods in R and evaluated the computational time of replicability analysis based on paired $p$-values. Computations were carried out in an Intel(R) Core(TM) i7-9750H 2.6Hz CPU with 64.0 GB RAM laptop. In the simulation studies, we set $\mu_1 = \mu_2 = 2.5, \sigma_1 = \sigma_2 = 1, \xi_{11} = 0.9, \xi_{01} = \xi_{10} = 0.025$, and $\xi_{11} = 0.05$. Let $m = 10,000, 20,000, 50,000$, and $100,000$, respectively. Table S1 summarizes the computational time of different methods to finish one replication with different numbers of genes. We observe that the computation is fast for all methods except MaRR [6] and IDR [5]. JUMP is scalable to hundreds of thousands of genes. The minor extra computational time of JUMP over other valid methods for replicability analysis can be ignored given its substantial power gain in replicability analysis. Table S2 summarizes the data information and computational time for replicability analysis on two pairs of SRT datasets from mouse olfactory bulb and mouse cerebellum.

Table S1: Computational time (in seconds) for replicability analysis in simulation studies

| # of genes | JUMP | BH | MaxP | radjust | MaRR | IDR | Šidák | Lancaster |
|---|---|---|---|---|---|---|---|---|
| 10,000 | 0.0280 | 0.0050 | 0.0040 | 0.0140 | 1.7564 | 4.1809 | 0.0040 | 0.0800 |
| 20,000 | 0.0530 | 0.0050 | 0.0050 | 0.0150 | 7.0752 | 8.9121 | 0.0050 | 0.1560 |
| 50,000 | 0.1280 | 0.0100 | 0.0080 | 0.0660 | 39.941 | 19.424 | 0.0070 | 0.3670 |
| 100,000 | 0.3230 | 0.0264 | 0.0200 | 0.0370 | 235.34 | 52.093 | 0.0170 | 0.7350 |

Table S2: Computational time (in seconds) for replicability analysis of different datasets: mouse olfactory bulb (MOB) and mouse cerebellum (MC).

| Dataset | # of genes/samples | | # of genes common to both studies | BH | MaxP | JUMP |
|---|---|---|---|---|---|---|
| | study 1 | study 2 | | | | |
| MOB | 9,547/237 | 10,680/1,185 | 8,547 | 0.0050 | 0.0040 | 0.0280 |
| MC | 17,481/14,667 | 20,117/11,626 | 16,519 | 0.0090 | 0.0070 | 0.0600 |

### References


[1] Y. Benjamini and Y. Hochberg. Controlling the false discovery rate: a practical and powerful approach to multiple testing. *Journal of the Royal Statistical Society: Series B (Methodological)*, 57(1):289–300, 1995.

[2] M. Bogomolov and R. Heller. Assessing replicability of findings across two studies of multiple features. *Biometrika*, 105(3):505–516, 2018.

[3] A. P. Dempster, N. M. Laird, and D. B. Rubin. Maximum likelihood from incomplete data via the em algorithm. *Journal of the royal statistical society: series B (methodological)*, 39(1):1–22, 1977.

[4] H. Lancaster. The combination of probabilities: an application of orthonormal functions. *Australian Journal of Statistics*, 3(1):20–33, 1961.

[5] Q. Li, J. B. Brown, H. Huang, and P. J. Bickel. Measuring reproducibility of high-throughput experiments. *Annals of Applied Statistics*, 5(3):1752–1779, 2011.

[6] D. Philtron, Y. Lyu, Q. Li, and D. Ghosh. Maximum rank reproducibility: a nonpara-

metric approach to assessing reproducibility in replicate experiments. *Journal of the American Statistical Association*, 113(523):1028–1039, 2018.

[7] Z. Šidák. Rectangular confidence regions for the means of multivariate normal distributions. *Journal of the American Statistical Association*, 62(318):626–633, 1967.

[8] P. L. Ståhl, F. Salmén, S. Vickovic, A. Lundmark, J. F. Navarro, J. Magnusson, S. Giacomello, M. Asp, J. O. Westholm, M. Huss, et al. Visualization and analysis of gene expression in tissue sections by spatial transcriptomics. *Science*, 353(6294):78–82, 2016.

[9] J. D. Storey and R. Tibshirani. Statistical significance for genomewide studies. *Proceedings of the National Academy of Sciences*, 100(16):9440–9445, 2003.

[10] S. Sun, J. Zhu, and X. Zhou. Statistical analysis of spatial expression patterns for spatially resolved transcriptomic studies. *Nature Methods*, 17(2):193–200, 2020.

[11] W. Sun and T. T. Cai. Oracle and adaptive compound decision rules for false discovery rate control. *Journal of the American Statistical Association*, 102(479):901–912, 2007.

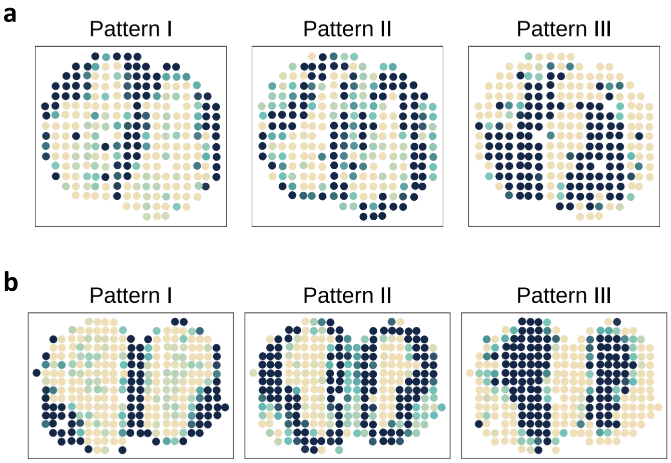


Figure S1: Spatial expression patterns summarized in two SRT studies on which the realistic simulation studies were performed. The SVGs were identified by SPARK [10] at an FDR level of $10^{-10}$. (a) Three spatial expression patterns based on 43 SVGs identified by SPARK [10] from Replicate 9 of the mouse olfactory bulb ST data. (b) Three spatial expression patterns based on 71 SVGs identified by SPARK from Replicate 12 of the mouse olfactory bulb ST data.

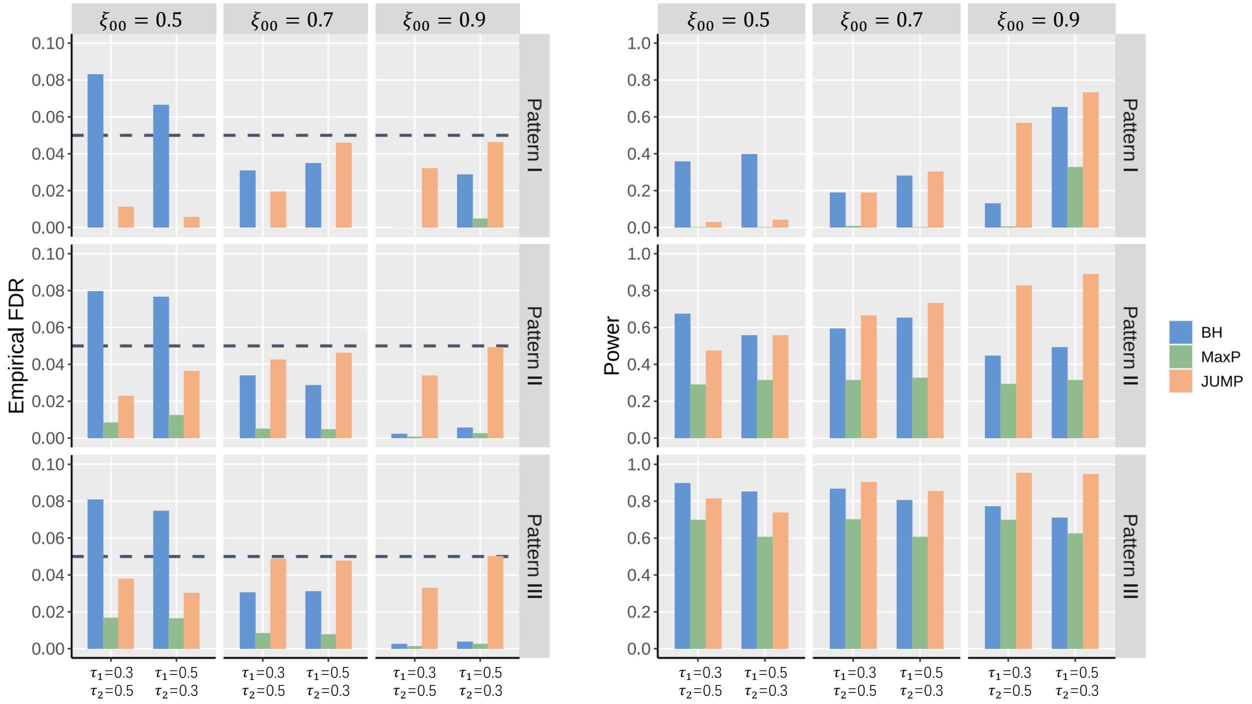


Figure S2: FDR control and power comparison of different methods in realistic simulations. Simulations were performed with $m = 10,000, \xi_{11} = 0.05$ and $\xi_{01} = \xi_{10}$. The signal strengths were set to be strong for study 1 and moderate for study 2. Each column corresponds to a different $\xi_{00}$ setting. Each row corresponds to a different spatial expression pattern on which the paired count data were generated. Patterns I-III for two studies are shown in Supplementary Fig. S1. In each panel, the empirical FDR and power of different methods were calculated at a target FDR level of 0.05 (horizontal dashed line in the plots) for different standard deviations (left: $s_1 = 0.3, s_2 = 0.5$; right: $s_1 = 0.5, s_2 = 0.3$).

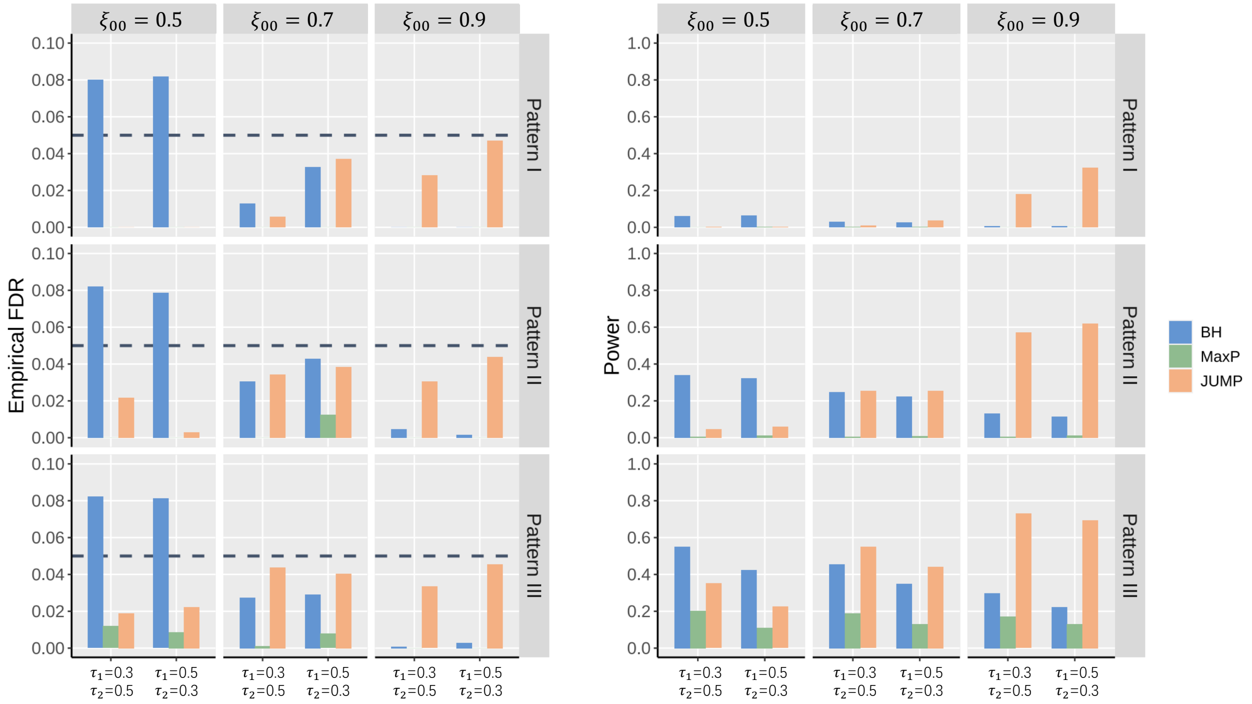


Figure S3: FDR control and power comparison of different methods in realistic simulations. Simulations were performed with $m = 10,000, \xi_{11} = 0.05$ and $\xi_{01} = \xi_{10}$. The signal strengths were set to be moderate for both studies. Each column corresponds to a different $\xi_{00}$ setting. Each row corresponds to a different spatial expression pattern on which the paired count data were generated. Patterns I-III for two studies are shown in Supplementary Fig. S1. In each panel, the empirical FDR and power of different methods were calculatedat a target FDR level of 0.05 (horizontal dashed line in the plots) for different standard deviations (left: $s_1 = 0.3, s_2 = 0.5$; right: $s_1 = 0.5, s_2 = 0.3$).

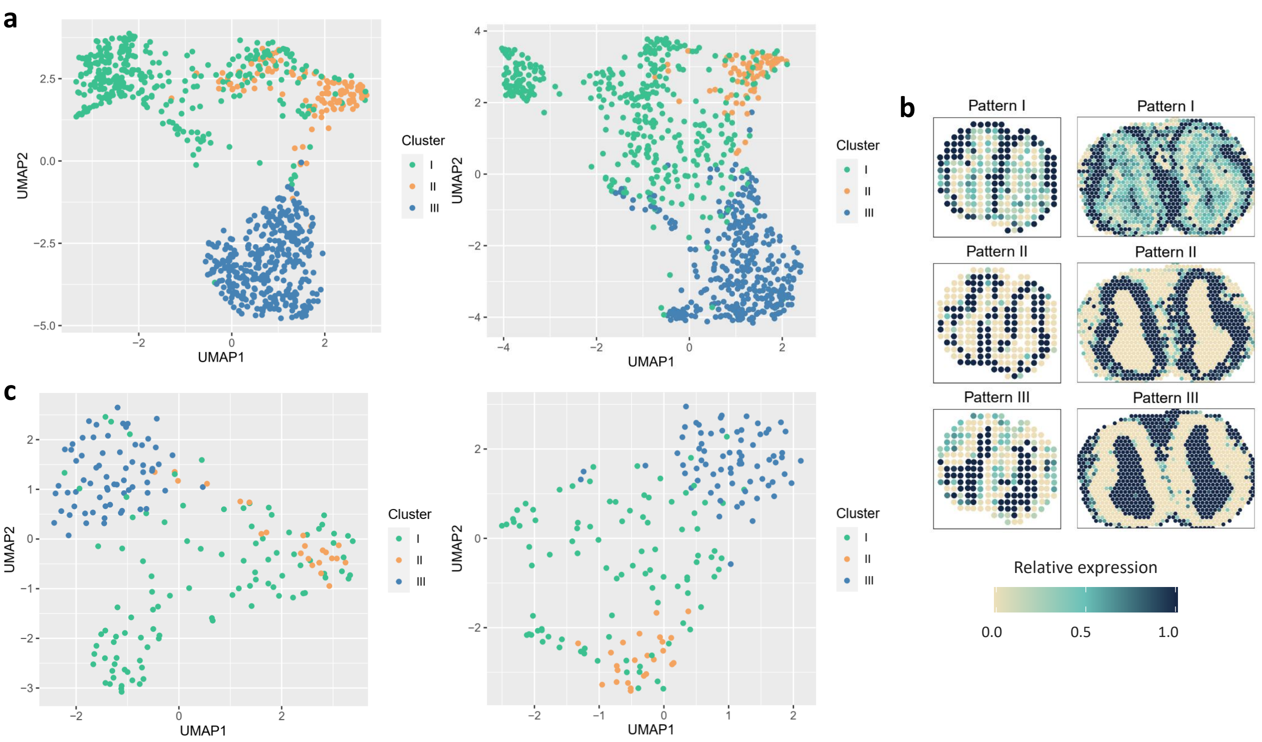


Figure S4: Analysis results of the mouse olfactory bulb data. (a) Scatter plot of 807 replicable SVGs identified by JUMP in two datasets (left: ST; right: 10X Visium). We first used UMAP (R package *umap*) to reduce the dimension to two. Then we used the cell labels obtained from hierarchical agglomerative clustering (R package *amap*) to visualize the distribution of cells in each cluster. (b) Spatial expression patterns in the two datasets summarized based on the 189 replicable SVGs additionally identified by JUMP (left: ST; right: 10X Visium). (c) Scatter plot of 189 replicable SVGs additionally identified by JUMP.

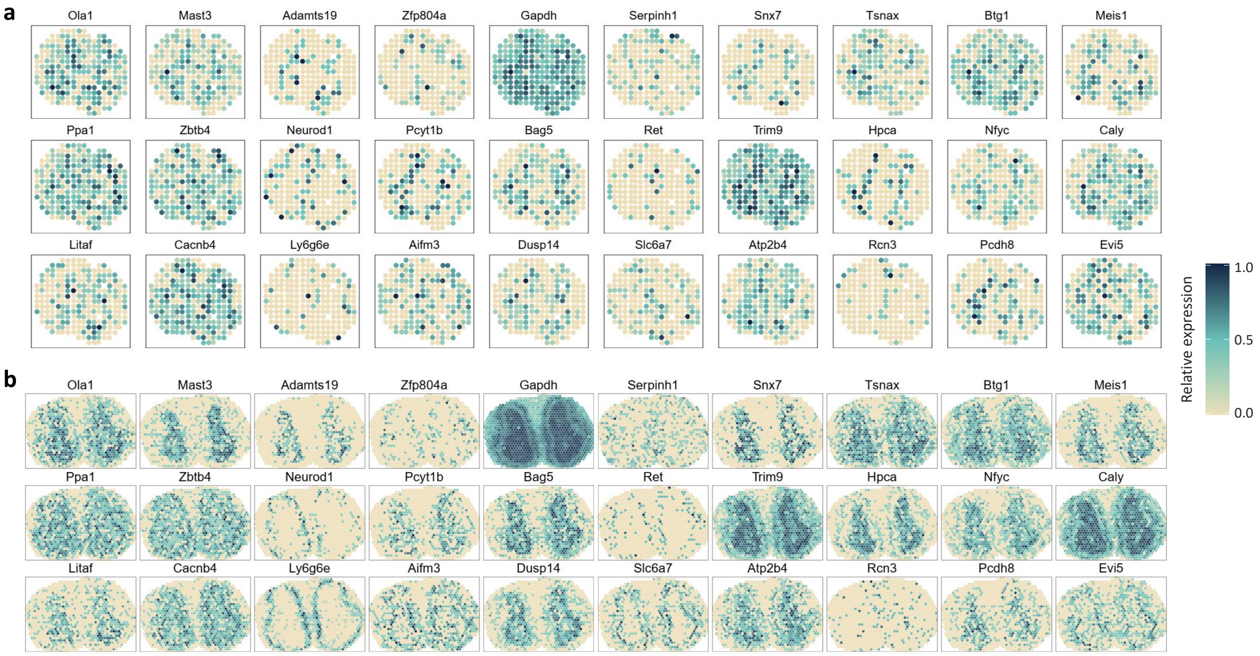


Figure S5: Spatial expression patterns of 30 genes randomly selected from the 189 replicable SVGs additionally identified by JUMP in mouse olfactory bulb. (a) Spatial expression patterns of the 30 randomly selected genes based on the ST data. (b) Spatial expression patterns of the 30 randomly selected genes based on the 10X Visium data.

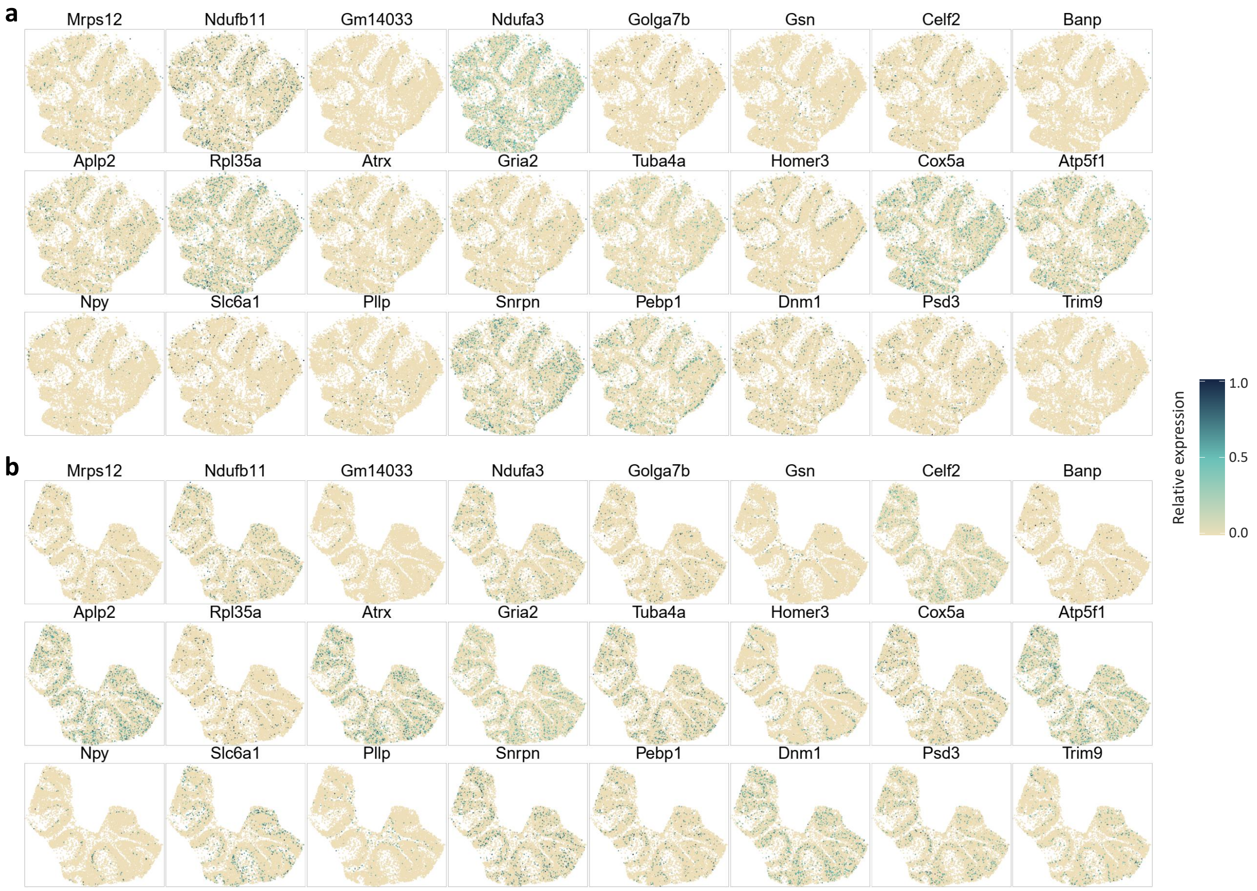


Figure S6: Spatial expression patterns of 24 genes randomly selected from the 169 replicable SVGs additionally identified by JUMP in mouse cerebellum. (a) Spatial expression patterns of the 24 randomly selected genes based on the Slide-seq data. (b) Spatial expression patterns of the 24 randomly selected genes based on the Slide-seqV2 data.
